# Likelihoods for a general class of ARGs under the SMC

**DOI:** 10.1101/2025.02.24.639977

**Authors:** Gertjan Bisschop, Jerome Kelleher, Peter Ralph

## Abstract

Ancestral recombination graphs (ARGs) are the focus of much ongoing research interest. Recent progress in inference has made ARG-based approaches feasible across of range of applications, and many new methods using inferred ARGs as input have appeared. This progress on the long-standing problem of ARG inference has proceeded in two distinct directions. First, the Bayesian inference of ARGs under the Sequentially Markov Coalescent (SMC), is now practical for tens-to-hundreds of samples. Second, approximate models and heuristics can now scale to sample sizes two to three orders of magnitude larger. Although these heuristic methods are reasonably accurate under many metrics, one significant drawback is that the ARGs they estimate do not have the topological properties required to compute a likelihood under models such as the SMC under present-day formulations. In particular, heuristic inference methods typically do not estimate precise details about recombination events, which are currently required to compute a likelihood. In this paper we present a backwards-time formulation of the SMC and derive a straightforward definition of the likelihood of a general class of ARG under this model. We show that this formulation does not require precise details of recombination events to be estimated, and is robust to the presence of polytomies. We discuss the possibilities for inference that this opens.

## Introduction

Following recent breakthroughs in simulation and inference methods, there is now strong interest in applying Ancestral Recombination Graphs (ARGs) to a variety of questions in evolutionary biology (Lewanski et al., 2024; Brandt et al., 2024; Nielsen et al., 2025). ARGs describe the intricately linked paths of genetic inheritance resulting from recombination (Hudson, 1983; Griffiths and Marjoram, 1996; Wong et al., 2024), and contain rich detail about ancestral processes. Numerous applications taking advantage of this detailed history in inferred ARGs are now appearing (Stern et al., 2019; Fan et al., 2022; Hejase et al., 2022; Guo et al., 2022; Ignatieva et al., 2022; Wang and Coop, 2022; Zhang et al., 2023; Nowbandegani et al., 2023; Ignatieva et al., 2023; Fan et al., 2023; Osmond and Coop, 2024; Huang et al., 2025; Grundler et al., 2024; Korfmann et al., 2024; Deraje et al., 2024; Speidel et al., 2025) and it seems likely that many more will follow (Harris, 2019, 2023). Although there are many different approaches to ARG inference (Wong et al., 2024), two broad classes of method have been the focus of most recent interest.

The first class of inference methods are Bayesian approaches that sample ARGs under a population genetics model such as the coalescent with recombination (Hudson, 1983), and its approximation, the Sequentially Markovian Coalescent (McVean and Cardin, 2005; Marjoram and Wall, 2006). ARGweaver (Rasmussen et al., 2014; Hubisz et al., 2020) has been the most widely used and studied, and until recently consistently out-performed other methods in terms of accuracy in a variety of benchmarks (Brandt et al., 2022), and has successfully been applied in several contexts (e.g., De Manuel et al., 2016; Shriner and Rotimi, 2018; Hejase et al., 2020; Stankowski et al., 2024). The recently-introduced method SINGER takes a similar approach, and by using an improved Monte Carlo algorithm promises to be even more accurate and substantially more scalable (Deng et al., 2024). The second class of inference methods that has been of recent interest are based on heuristics and approximate models. Relate (Speidel et al., 2019), tsinfer (Kelleher et al., 2019), ARG-Needle (Zhang et al., 2023) and Threads (Gunnarsson et al., 2024) work on quite different principles, but share some common properties. Firstly, they are all heuristic and approximation driven, favouring computational scalability over a basis in a statistical model. Secondly, they all produce a single ARG as output, inferring a deterministic point estimate. Thirdly, they regard the precise timing of nodes as a separate problem, focusing only on the relative ordering of nodes when producing the initial ARG. General purpose ARG dating methods are appearing (Wohns et al., 2022; Deng et al., 2025) and can also improve dating performance in ARGs sampled from the SMC (Deng et al., 2025). Fourthly (and most importantly for the purposes of this paper) the ARGs that heuristic methods produce lack some of the topological information present in the output of ARGweaver or SINGER. This occurs, roughly speaking, because of a basic difference in their approach to uncertainty.

There is considerable uncertainty in ARG inference, and often a fundamental lack of information in the sequence data to distinguish different possibilities. Consider, for example, a case in which we have three ancestral lineages with identical sequences. We know that there must have been two coalescence events, but there is no mutational information to distinguish their relative ordering. Sampling methods overcome this problem by randomly choosing an ordering, with the uncertainty communicated by the order changing in the different ARGs sampled from the posterior. The other approach (used by methods such as tsinfer) is to communicate the uncertainty structurally by means of a polytomy: we have no information about the intermediate coalescent event, and so we omit it. Similarly, there is often a fundamental lack of information about the details of recombination events, and heuristic methods (in different ways) omit these details. In many ways, the significant progress made in scalability by these heuristic methods is *because* they don’t attempt to precisely reconstruct recombination events.

There are substantial advantages to the statistically rigorous sampling approaches of ARGweaver and SINGER. In particular, they tend to be more accurate (at the scale at which they can be compared), and sampling ARGs from a posterior distribution provides a means of quantifying the uncertainty around estimates. On the other hand, heuristic methods have the potential to use the greater amount of information in large datasets, capturing fine-scale details about the recent past. Although SINGER is a major step forward in terms of scalability over ARGweaver and other methods, inference is still only feasible for hundreds of samples. Meanwhile, tsinfer, ARG-Needle and Threads have been applied to datasets of hundreds of *thousands* of samples. With the explosive growth in dataset sizes in recent years (Caulfield et al., 2017; Turnbull et al., 2018; Bycroft et al., 2018; Backman et al., 2021; Halldorsson et al., 2022; UK Biobank Whole-Genome Sequencing Consortium et al., 2023; All of Us Research Program Genomics Investigators et al., 2024; Stark et al., 2024; Cook et al., 2025) there is a pressing need for more statistical rigour in large scale ARG inference.

A key problem facing the field is that the ARGs estimated by heuristic large-scale methods lack a concrete connection to population genetic theory because we cannot compute their likelihood under the SMC or similar models. Computing the likelihood of an ARG under a model is a fundamental element of probabilistic inference. The classical Kuhner-Yamato-Felsenstein (2000) formulation (henceforth: KYF) works by assigning a probability to each recombination and common ancestor event, and to the inter-event waiting times. Although the KYF formulation is a natural and elegant way to describe the likelihood of an ARG, the high level of detail about the underlying events that is required is not present in many estimated ARGs (Wong et al., 2024) or even in most simulations. Simulators such as the classical ms program (Hudson, 2002) output ARGs in a tree-by-tree format, which does not contain sufficient detail for likelihood computation. The msprime simulator (Kelleher et al., 2016; Baumdicker et al., 2022) by default outputs a more complete and efficient representation of the simulated ancestry, but still does not provide the exhaustive detail required. Indeed, providing the information required to support likelihood calculations originally motivated the addition of the `record_full_arg` option to msprime (Baumdicker et al., 2022). Forwards-time simulations that output ARGs (Kelleher et al., 2018; Haller et al., 2018) also do not typically contain the level of detail required to compute a likelihood under the KYF approach.

In this paper we provide a new formulation of ARG likelihood which has much more lenient input requirements than KYF, and gracefully handles incomplete and imprecise ARGs. To do this, we give a backwards-in-time description of the SMC which naturally leads to a straightforward method for computing the likelihood of any ARG, and describe an efficient algorithm to do so. We show that explicitly identifying recombination events via nodes in the graph is not necessary (from the perspective of computing the likelihood) as long as the “locally unary” (Wong et al., 2024; Fritze et al., 2024) sections of common ancestor nodes are retained. We also show via an empirical example that the likelihood is robust to the presence of polytomies and other imprecise information, and illustrate how it can be decomposed into separately evaluated time slices. Finally, we discuss how these results may be applied and extended.

## Results

### ARGs

Although the term Ancestral Recombination Graph (ARG) has been used to mean several things, in the generic sense, it is a labelled graph structure that records the inheritance of genetic material (Wong et al., 2024). An ARG of the sort we consider describes the history of inheritance of the genomic material of a focal set of individuals, or *samples*. Concretely, an ARG is equivalent to a collection of (non-contradictory) statements of the form “*c* inherited from *p* on the genomic segment from *x* to *y*”, where *c* and *p* are (haploid) genomes, and *x* and *y* are genetic coordinates. We record each such statement in an “edge”, often summarised as the tuple (*p, c, x, y*). By “describes the history”, we mean that each node in the ARG represents a specific (ancestral) genome, and that the ARG specifies the genealogical tree describing how the samples are related at each point on the genome.

An ARG can therefore be thought of either as a collection of inheritance relationships between ancestral haplotypes, or as a sequence of genealogical trees that have common node labels. Fig.1 shows a small ARG that describes relationships between four samples on a 100bp piece of genome, represented as both a graph (top) and a series of local trees (bottom). Both structures represent the same information: for instance, the fact that the ancestral genome at *A* inherited the left half of its genome from ancestor *g* and the right half from ancestor *f* is seen in the fact that the first two trees have *A* below *g*, and the remaining have *A* below *f*. The blue dotted line from *A* in the second tree to *f* in the third tree draws our attention to the fact that the path traced by *A*’s ancestry switches at that point. In fact, *A* is in blue because it is a *recombinant* node – an ancestor that has a segment of genome inherited by a sample, but that segment was inherited in two pieces from their parent. Other nodes (labelled by black numbers) are either samples or *common ancestry* nodes – ancestors that are the most recent common ancestor for two or more samples at some place along the genome.

**Figure 1:**
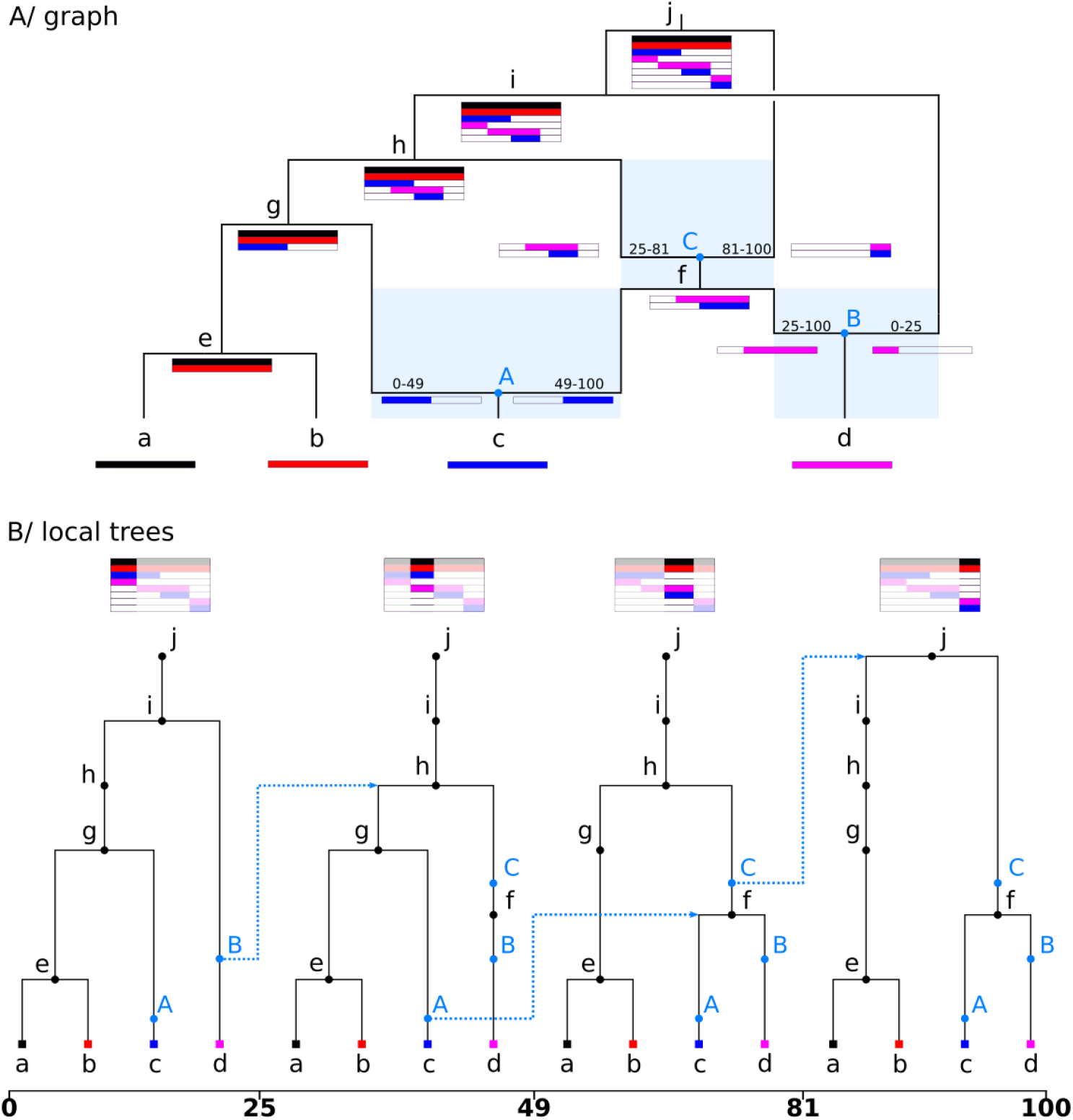
An ARG generated under the SMC, represented as a graph (top) and as a series of local trees (bottom). To ensure the one-to-one correspondence between both representations a node is recorded for each local tree whenever a node is encountered along its ancestry path in the graph representation. This results in nodes that (locally) have only a single child, and are therefore (locally) unary. Nodes A, B and C are recombinants, i.e, have two parents in the graph and are unary in the local trees. The blue shaded regions indicate the range of possible times associated with these recombinants. The blue arrows show the standard left-to-right logic of the SMC whereby the floating lineage coalesces randomly with a the remaining portion of the local tree following recombination.

Crucial for the equivalence between the two representations is the presence of nodes that only have a single child along one or more local trees – i.e., are *locally unary* (Wong et al., 2024). Recombinant nodes (in blue) have a single child in all trees. However, common ancestry nodes can be locally unary as well: for instance, sample *d* inherited a longer stretch of genome from ancestor *f* than sample *c*. In the top plot, this is visible because the pink bar below node *f* extends further to the left than does the blue bar, while in the bottom plot, this is visible because node *f* is parent to *d* in the second tree, but not to node *c*, and so is unary.

These locally unary nodes enable us to uniquely identify all lineages that were hit by a recombination event, even if we simplify the ARG by removing all recombinant nodes (*A, B*, and *C* in Fig.1). In the absence of these recombinant nodes, a recombination can be observed easily enough as a change in parent going from one genomic region to the next. Without the recombinant nodes we no longer know the precise time of each recombination event, but we do still have constraints on it; for instance, we know that *C* happened somewhere between node *f* and node *h*. More generally, a recombinant node must happen somewhere between the child node and the most recent of the parent nodes. This is again easier to visualise using the graph representation (blue shaded area, Fig.1). Although these unary nodes are useful and to some degree clearly inferrable, they are not always represented by simulators or ARG inference methods. However, to our understanding this is because the field is used to thinking in terms of the trees rather than in terms of relationships between haplotypes, not because of any question of knowability. See Fritze et al. (2024) for more discussion on unary nodes and haplotype aware ARG comparison metrics.

### SMC backwards-in-time

The coalescent process describes the ancestry of a set of sampled genomes. The original algorithm to simulate the coalescent with recombination was formulated backwards in time by Hudson (1983). Wiuf and Hein later described a tree-by-tree method to simulate the same stochastic process (Wiuf and Hein, 1999b). This along-the-genome approach is considerably more complex as the next tree to be simulated depends on all previous trees. This idea however provided the basis of the SMC (McVean and Cardin, 2005), restricting the set of all state space transitions possible under the Hudson coalescent to obtain a process that is Markovian both backwards-in-time, as well as along-the-genome. In its backwards-in-time formulation, the SMC only requires a simple modification to the coalescent with recombination. Instead of allowing any pair of lineages to coalesce, the SMC is the process in which only lineages with *overlapping* genomic intervals may coalesce. The SMC’ is a refinement of the SMC which also allows lineages with abutting genomic intervals to coalesce (Marjoram and Wall, 2006). Although McVean and Cardin (2005) verbally described the SMC using this backwards-in-time description, their analysis was almost entirely in left-to-right terms, and subsequent papers followed this trend (Li and Durbin, 2011; Paul et al., 2011; Schiffels and Durbin, 2014; Rasmussen et al., 2014).

Linking the resulting tree-valued process along the genome back to the relationship between the “graph” and “local trees” representation of ARGs, the Markovian property can be formulated as follows. The distribution of the next tree within the series of local trees can be generated only knowing the current tree, provided that we remove recombinant nodes entirely, and erase the unary portions of coalescent nodes. To see that the locally unary nodes make the process non-Markov, consider that a node in a tree might be unary either because it’s *about to be* (in the left-to-right sense) a coalescent node or it *already has been*: looking at the second tree in Fig.1, we know node *f* will be coalescent further along the sequence because it was not present in the first tree. So, the first tree gives us information about subsequent trees that the second tree alone could not. Further note that by limiting common ancestry events to overlapping pairs of lineages, we are guaranteed that all nodes in the ARG only carry ancestral material, that is, genetic material that is inherited by either one of a set of samples (Wiuf and Hein, 1999a). We therefore require a single half-open interval to describe the ancestral material associated with each node in the ARG.

We now define the process more formally, by reintroducing the notation of McVean and Cardin (2005) required to describe the likelihood computation. This section does not deviate from the original description of the SMC apart from how coalescable pairs are counted. This subtle nuance greatly simplifies the ability to keep track of coalescable pairs going backwards in time. At any point in time, the state of the process is the set of labelled lineages extant at time *t, L*(*t*) = {*X*_*j*_(*t*)}. A lineage *X*_*j*_(*t*) = [*x*_*j*_, *y*_*j*_) is labelled by an integer *j* and defines the half-open genomic interval of its ancestral material. Because we look backwards in time, *t* = 0 is today, and *L*(0) consists of *n* sampled lineages, labelled 0 to *n −* 1, each represented by a single interval spanning the entire genome. Then, the process evolves by a succession of common ancestor and recombination events until each segment of ancestral material is only present in one lineage. The waiting time to next event is determined by these two competing processes with exponentially distributed waiting times as outlined below.

Recombination is described by a Poisson process of rate *r* per unit of genomic length and time: so, if *T* (*t*) = ∑_*j*_ |*X*_*j*_(*t*)| is total length of ancestral material carried by lineages at time *t*, then recombination occurs at instantaneous rate *rT* (*t*). A recombination to the left of *x* (i.e., between base pairs *x* and *x −* 1) with *x*_*j*_ *< x < y*_*j*_ splits lineage *j* into two new lineages, given new, unique labels. We then remove lineage *j* and add the two new lineages to the state *L*(*t*). This operation keeps the total amount of ancestral material unchanged.

Assuming a randomly mating, diploid population of constant size *N*_*e*_, common ancestry occurs between two *overlapping* lineages at rate *λ* = 1*/*(2*N*_*e*_). Here we deviate from the McVean and Cardin formulation, and define a strict total order on all lineages in *L*(*t*): for each lineage *X*_*j*_(*t*) = [*x*_*j*_, *y*_*j*_) we define its order by the left coordinate *x*_*j*_ and label *j* (to break ties). The instantaneous rate of coalescence then equals *λ* multiplied by the number of overlapping pairs, i.e., 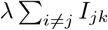, where

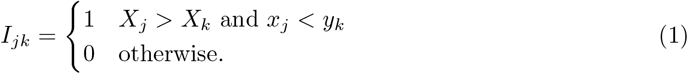

We say that lineage *k* can coalesce *into* lineage *j* if *I*_*jk*_ = 1. Note that since *X*_*j*_ *> X*_*k*_ implies that *x*_*j*_ *≥ x*_*k*_, this condition implies that the left endpoint *x*_*j*_ of *X*_*j*_ falls in the interval defined by *X*_*k*_. From the definition of the strict total order it follows that if *I*_*jk*_ = 1 then *I*_*kj*_ = 0. The newly formed lineage acquires the union of both intervals, the original lineages are removed, and *L*(*t*) is updated accordingly. For later use, also define *C*_*j*_(*t*) to be the set of lineages that can coalesce into lineage *j*, i.e., *C*_*j*_(*t*) = *{X*_*k*_ *∈ L*(*t*)|*I*_*jk*_(*t*) = 1*}*; thus, |*C*_*j*_(*t*)| = *I*_*j*_(*t*) = ∑_*k*_ *I*_*jk*_(*t*), and the total rate of coalescence at time *t* is ∑_*j*_ |*C*_*j*_(*t*)|. Because *I*_*jk*_ is defined in terms of a total order, the sets *C*_*j*_(*t*) are disjoint, which will simplify likelihood computations later. In particular, although recombination affects *which* lineages can coalesce into *j*, it does not change the *number* of such lineages: in other words, a recombination event changes *C*_*j*_(*t*) but not *I*_*j*_(*t*).

### ARG likelihood

We have defined the SMC process in terms of rates of various types of events. In general, the likelihood for a continuous-time Markov process specified in this way is exp(*−*Λ) ∏_*i*_ *λ*_*i*_, where *λ*_*i*_ is the rate of the *i*^th^ realised event, and Λ is the sum of all rates of all possible events integrated over the entire process (also called the “total hazard”). As discussed in the Introduction, in many cases these events are not what is estimated by inference methods, which produce, more fundamentally, a collection of genetic inheritance relationships (edges) between ancestral haplotypes (nodes). Our goal, therefore, is to decompose the overall likelihood into the per-node and edge contributions, allowing us to compute likelihoods for the incomplete ancestries estimated by current methods. Using the out- and in-degree of the nodes we’ll first reason about the number of common ancestor events and recombination events that could have resulted in the observed ARG. Next the information contained in the edges is leveraged to compute the total hazard associated with both types of events.

Let deg_*o*_(*u*) = |*{e ∈ E* : *p*_*e*_ = *u}*| be the out-degree of a node *u*, i.e., the number of edges having *u* as a parent. Because mergers are binary in the SMC, this must have been the result of deg(*u*) *−* 1 common ancestry events, and so this contributes a factor of 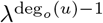 to the likelihood for nodes with any children. Note however that this description is general and holds for the case of more-than-binary mergers. By treating such polytomies as if there were zero-length edges between a sequence of mergers, the order of these unresolved mergers does not affect the likelihood. Under the SMC the likelihood of a zero-length edge is zero. This is not special: since edge lengths are continuously distributed, the likelihood of any particular value is also zero.

Similarly, let deg_*i*_(*u*) = |*{e ∈ E* : *c*_*e*_ = *u}*| be the in-degree of a node *u*, i.e., the number of edges having *u* as a child. Each additional edge that is ancestral to a given node implies one additional ancestral recombination event, and thus a factor of 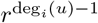 in the likelihood, if the node has any parents. Put together, these two contribute

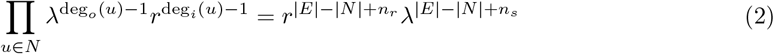

to the likelihood, where *n*_*r*_ is the number of nodes with in-degree 0 (roots), and *n*_*s*_ is the number of nodes with out-degree 0 (usually, samples). (The simple form follows because each edge contributes exactly one to some node’s out-degree and some node’s in-degree.)

Now consider the “hazard” associated with recombination. Begin with an edge *e* = (*p, c, x, y*). This implies that there was a sequence of lineages from the time of the child *c* (call this time *t*_*c*_) back to the time of the parent *p* (again, *t*_*p*_), that contained the segment [*x, y*), and the lineage that directly inherited from *p* had the segment [*x, y*) (see Fig. 2). We therefore know there was no recombination between *x* and *y* on any of those lineages over this time span, the probability of which is exp(*−r𝒜*_*e*_), where *r* is the recombination rate and *𝒜*_*e*_ is the total area (eligible links multiplied by length in time) of the edge. The length in time of the edge is just *t*_*p*_ *− t*_*c*_, while the number of “eligible links” in the edge is the number of adjacent base pairs in the edge on which recombination did not occur, which is either *y − x* (if this edge is the leftmost of all edges above node *c*) or *y − x −* 1 (otherwise). So, the first contribution to the likelihood from this edge *e* is

**Figure 2:**
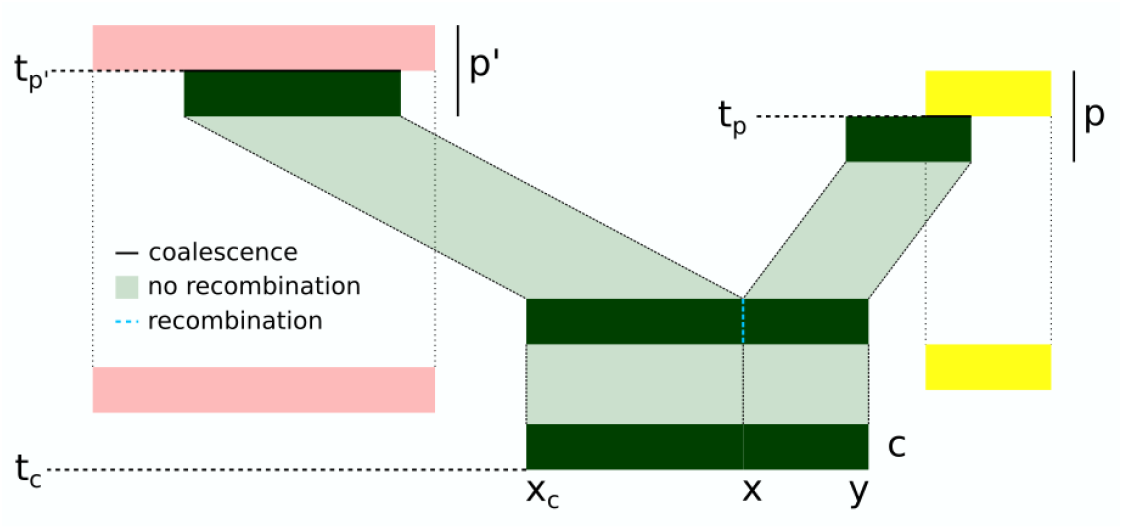
Graphical representation of the events associated with the edge (*p, c, x, y*). The lineage carried by *c* (dark green bar) spans [*x*_*c*_, *y*) until a recombination event (blue dotted line) splits it into two segments. Left and right hand segments coalesce (with the pink and yellow lineages, respectively) at times *t*_*p*_*′* and *t*_*p*_ respectively, bounding the time to recombination by min(*t*_*p*_, *t*_*p*_*′*).

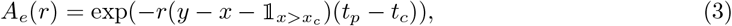

where as for lineages, *x*_*c*_ is the leftmost position of the edges whose children are *c*.

Finally, we turn to the hazard associated with common ancestry events. To do this, we need only consider what happens along the left side of each edge. Following what we did for lineages, we define a total order on edges by ordering by left endpoint and breaking ties by child (i.e., (*p, c, x, y*) *<* (*p*^*′*^, *c*^*′*^, *x*^*′c*^, *y*^*′*^) if *x < x*^*′*^, or if *x* = *x*^*′*^ and *c < c*^*′*^). The instantaneous rate of coalescence of a lineage whose left endpoint is at *x* at a given time is equal to *λ* multiplied by the number of earlier lineages that overlap *x* at that time. Using this total order, we can define *I*_*e*_(*t*) to be the total number of earlier edges that overlap edge *e* and are present at time *t*. Then, the cumulative hazard for common ancestry events of edge *e* between time *s* and *t* is given by 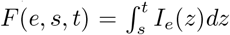. If the edge *e* = (*p, c, x, y*) is the leftmost segment of material inherited by node *c* (e.g., the left hand segment ancestral to node *c* in Fig. 2), then no recombination occurred along this edge and this edge simply contributes exp(*−F* (*e, t*_*c*_, *t*_*p*_)) to the likelihood. However, if the edge is *not* the leftmost segment (as shown in Fig. 2), then it was initiated by a new recombination breakpoint, and we need to integrate over possible times the breakpoint occurred. If this is the case, there is another edge (*p*^*′*^, *c, x*^*′*^, *x*) with the same child node, immediately adjacent to the focal edge (*p, c, x, y*). The recombination event that split the two must have happened between *t*_*c*_ and the smaller of *t*_*p*_ and *t*_*p*_*′*. Taking all this into consideration, the contribution to the likelihood here is

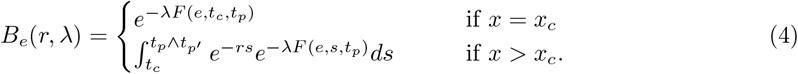

The overall likelihood of an ARG *𝒢* defined by the set of edges *E* and nodes *N* given parameters *θ* is then given by

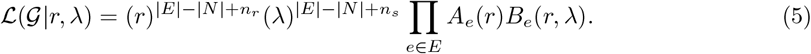

One significant advantage of this formulation is that the computation time is linear in the number of edges, and an efficient algorithm exists to keep track of *I*_*e*_(*t*) (see Appendix). Note that we have only considered the topology and branch length information here and not considered mutational information. Assuming the infinite sites model, computing the likelihood of an ARG given a mutation rate *µ* and set of mutations is straightforward, as the overall likelihood can be directly decomposed into the per-edge contributions (Baumdicker et al., 2022; Mahmoudi et al., 2022).

### Robustness to partial and incorrect information

Eq. (5) allows us to compute a likelihood for *any* collection of nodes (ancestral haplotypes that lived at some time *t* in the past) and edges (a record that child node *c* inherited the genomic interval [*x, y*) from parent node *p*) under the SMC. Other than some basic consistency constraints (e.g., parents must be older than children) there are no requirements on the topology described, and in particular, no requirement that the events from the model (the SMC) should be explicitly identified in the graph. Our formulation reasons about the events that *could have resulted in* the observed nodes and edges, and therefore frees us from the requirement of having to explicitly write those down. In this section we illustrate the robustness of this approach to incomplete and potentially incorrect information about recombination and coalescent events via a proof-of-concept example.

In Fig 3 we begin with a realisation of the SMC containing complete information about every simulated event (Full ARG; 9,378 nodes; 13,221 edges; 2,933 trees). We then plot the likelihood computed by Eq. (5) as we vary the recombination rate parameter *r*, as a ratio of the likelihood at the true value of *r*. Reassuringly, we can see that the minimum of the likelihood-ratio curve when using the full simulated ARG is close to the true value of the parameter. (The likelihood ratio curve obtained when computing the likelihood under the full coalescent with recombination model using the KYF formula is identical; not shown.) We next compute the likelihood ratio for the ARG obtained when we simplify out the recombination nodes from the full ARG (“RE nodes removed”; 3,332 nodes; 6,901 edges; 2,933 trees; note that msprime uses distinct nodes for the left and right parent of each recombination). The curve we obtain is indistinguishable from that of the full ARG, demonstrating that the locally unary portions of coalescent nodes contain all the information that we need about recombination events for this application. When we fully simplify the simulated ARG to remove all locally unary portions of nodes from trees (“Fully simplified”; 3,332 nodes; 10,674 edges, 2,933 trees) we can see there is a significant loss of information, and the minimum of the likelihood ratio surface is biased away from its true value, as expected, because removing the unary portions of nodes has removed some of the area of the ARG in which recombination did not occur. (See Wong et al. (2024) for more discussion on varying degrees of ARG simplification.)

**Figure 3:**
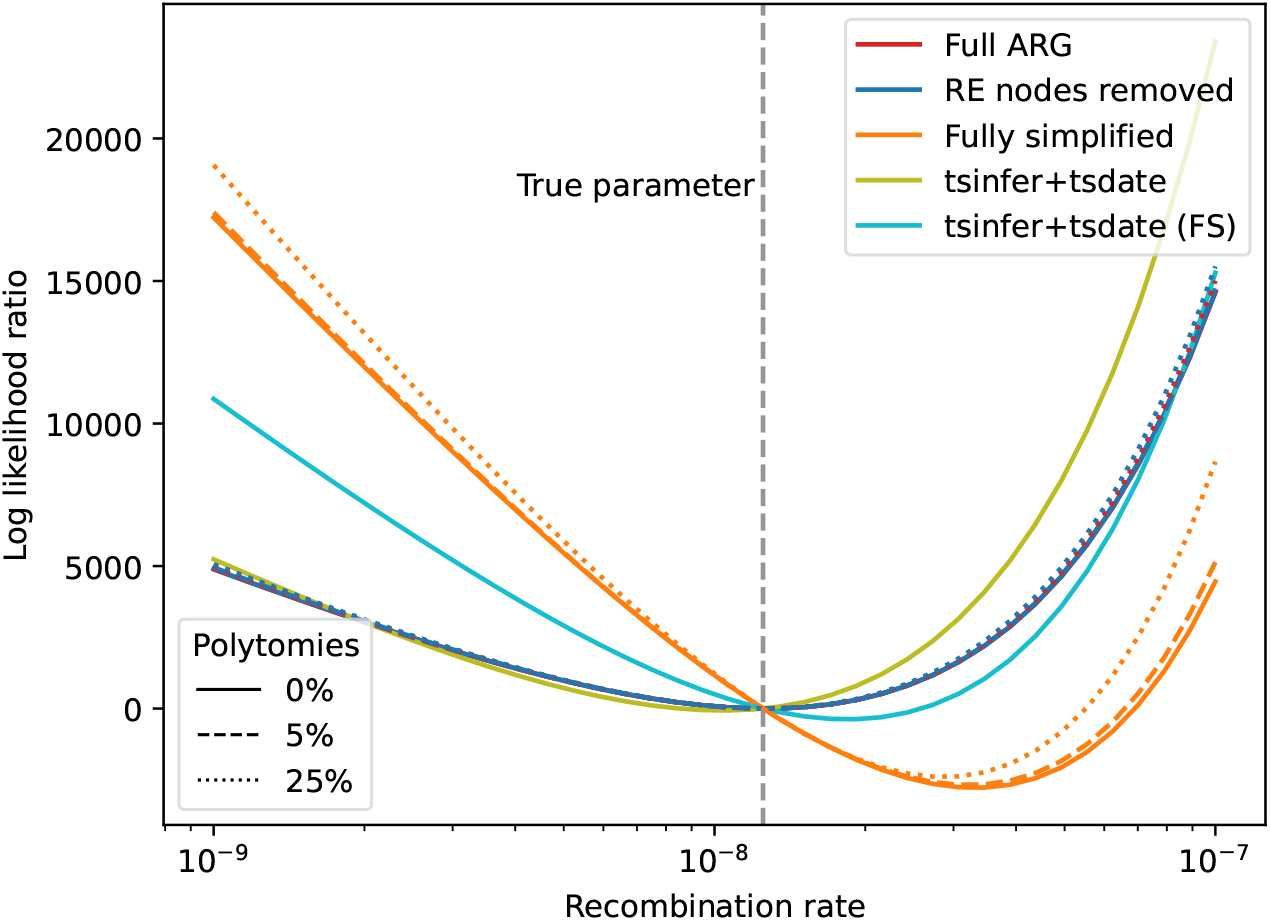
Log likelihood-ratio curves for ARGs with various properties. The “Full ARG” here is the result of an msprime simulation of the SMC (100 diploid samples, genome length 1Mb, *r* = *µ* = 2.5 *×* 10^*−*8^, *N*_*e*_ = 1 *×* 10^4^). The “RE nodes removed” ARG is the result of simplifying out all of the recombination nodes, and the “Fully simplified” ARG has all locally unary nodes removed. Each of these simulated ARGs additionally has 5% and 25% of the internal nodes removed to create polytomies. We also show the results for ARGs inferred from the original simulation data using tsinfer and tsdate (see text for details), in which we use the default output of tsinfer including unary nodes (tsinfer+tsdate), and in which we fully simplify before passing to tsdate (tsinfer+tsdate FS). See the text for details on how the likelihood ratio values are computed. The “Full ARG” and “RE nodes removed” lines coincide.

Fig 3 shows that Eq. (5) is also remarkably robust to the presences of polytomies in this example. For each of the three ARGs discussed in the previous paragraph, we also evaluated the likelihood for ARGs in which 5% or 25% of the internal ARG nodes are removed and edges adjusted accordingly. This introduces polytomies at varying degrees in the different ARGs because deleting a node in a tree (and connecting its children directly to its parent) will only create a polytomy if the parent is not unary. Thus, while deleting 25% of the internal nodes in the fully simplified ARG created an average of 33 polytomies over the 2,933 trees along the sequence (each tree having 200 leaf nodes), it resulted in an average of 21 polytomies per tree in the ARG where recombination nodes have been removed, and only 10 in the full ARG (which has an average of 1026 unary nodes per tree). Given this disparity in the number of polytomies introduced, the effects on the likelihood across the three examples are not entirely comparable. For the ARGs containing unary nodes, removing 25% of the internal nodes has negligible effect on the likelihood curve in Fig 3, and even on the fully simplified ARG has only a very minor effect on location of the minimum.

Fig 3 also shows results for ARGs inferred from the simulated data using tsinfer and ts-date (Wohns et al., 2022), using default parameters for tsinfer, and the true values of mutation rate and *N*_*e*_ for tsdate. In the first, we use the default output of tsinfer (2,148 nodes; 5,589 edges; 1,307 trees; 3,039 mutations). This ARG has an average of 192 unary nodes, 120 binary nodes, and 32 nodes with three or more children per tree. Tsinfer does not directly estimate recombination events; the breakpoints between trees are the consequence of switches between parental haplotypes in the Li and Stephens (2003) copying process. Information about recombination events in the returned ARG is therefore quite imprecise, and likely to contain incorrect and contradictory details if one were to attempt to reconstruct the exact subtree-prune-and-regraft moves (e.g., Rasmussen and Guo, 2023). Nonetheless, in this toy example at least, our likelihood function performs well on the output of tsinfer, with the minimum close to the true parameter value. We also show the likelihood curve obtained when we fully simplify the output of tsinfer (1,865 nodes; 7,511 edges; 1,252 trees; 3,039 mutations) before dating. We can see that, as with the true data, removing all the unary nodes results in a significant bias away from the true value of the recombination rate parameter.

### Piecewise decomposition over time

Our final example explores decomposition of the likelihood into separate components for different segments of time. By explicitly associating a single likelihood with each edge, we can compute the likelihood for any slice of the ARG taken in time and/or along the genome. In the case of a fully precise ARG, in which each recombination event has an exact time, this is true as long as the slicing operation does not erase the information on whether an edge is associated with the left or right hand segment generated by a recombination event. In the absence of the exact time to recombination, the integral in Eq. (4) means that the likelihood for the entire ARG does not factorise into the likelihoods of these individual slices. More precisely, the likelihood associated with any edge that is initiated by a recombination breakpoint cannot be factorised into the product of two integrals of the same form. However, as long as the individual slices of the ARG contain sufficient information we can approximate the true integral as the product of the integral of the slices. Fig. 4 demonstrates this idea on a toy example. We simulated an ARG under human-like parameters with two effective population size changes, and subsequently computed the posterior distribution on the three effective population sizes using a uniform prior on each slice independently. The result is reasonably accurate, despite only using 10kb of sequence.

**Figure 4:**
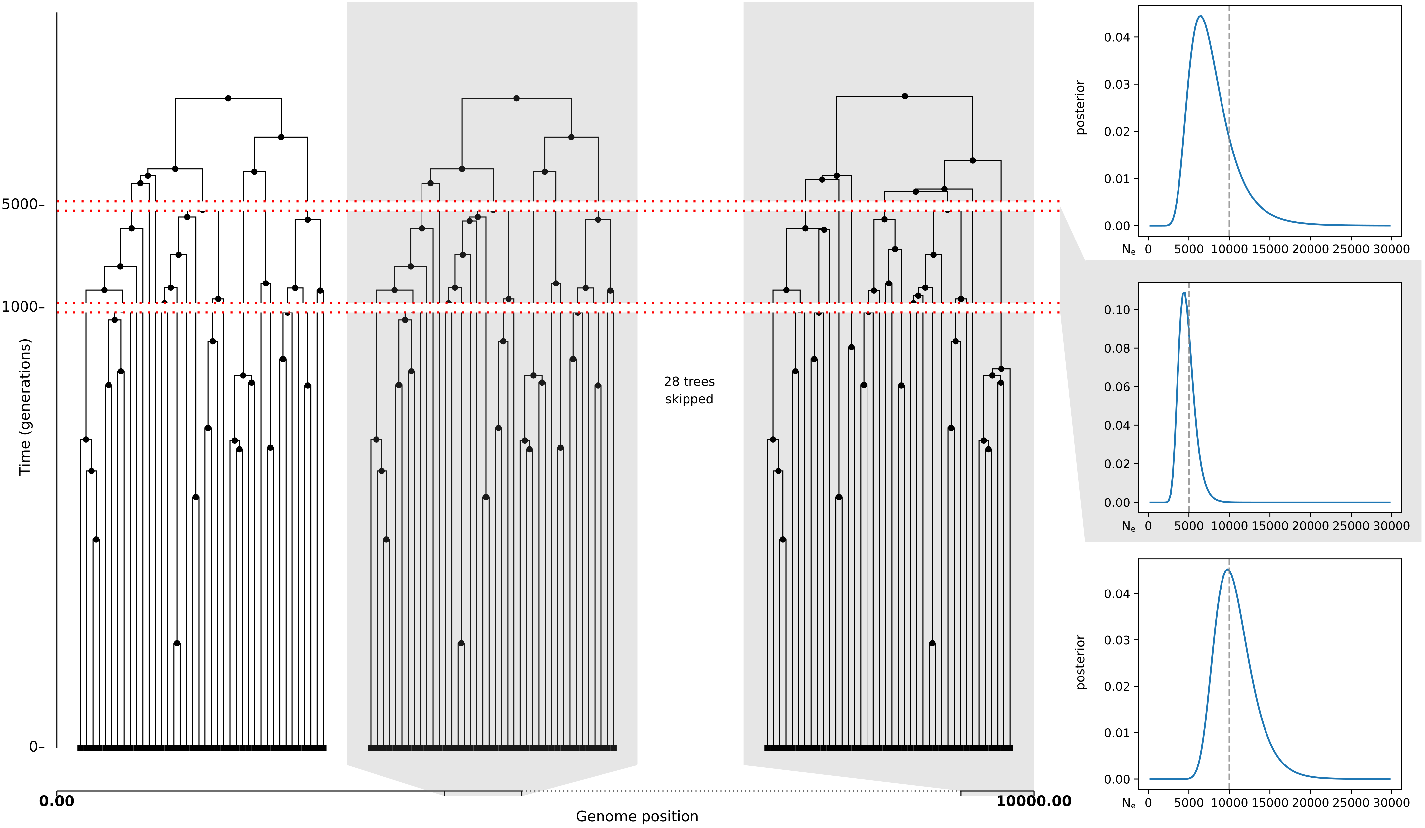
Simulated ARG under the Hudson coalescent (left). *N*_*e*_ is constant for each of the 3 time slices: from 0 to 1,000 generations ago *N*_*e*_ is 10,000; from 1,000 to 5,000 generations ago it is 5,000; and past that it is 10,000. Remaining parameters: genome length is 10kb, 20 diploid samples, and *r* = 2.5 *×* 10^*−*8^. There are 148 edges in the resulting ARG. A single *N*_*e*_ value was estimated for each of the three slices of the ARG separated by the true population size change time points. The inferred posterior distribution (right) using the SMC-likelihood was obtained using a uniform prior. The number of edges per time slice (from most recent to oldest) was: 68, 57, and 23. Locally unary nodes were omitted for clarity.

## Discussion

The key contribution of this paper is to rederive a classical ARG likelihood function in a way that supports the sort of partial and imprecise information provided by some modern ARG inference tools, using a novel backwards-time derivation of the SMC. The underlying process that we reason about probabilistically is still composed of events affecting lineages, but, because we do not necessarily observe or estimate these events, we integrate over the possible timings implied by the information in the input ARG. Our method can compute a likelihood under the SMC for a very general class of ARG (Wong et al., 2024), a significant increase in flexibility over the strict input requirements for the classical Kuhner-Yamato-Felsenstein formulation. Locally unary nodes play a key role, and although not currently a common focus of ARG research (Wong et al., 2024), they are estimated by tsinfer and may be imputed to a high degree of accuracy from fully simplified ARGs (Fritze et al., 2024). We have shown via some illustrative examples that the formulation is robust to incomplete and imprecise information, tolerating the loss of a significant fraction of the internal nodes in the ARG to minimal effect. We have also shown that the likelihood function performs well (at least in our toy example) on ARGs inferred by tsinfer+tsdate, successfully capturing information about latent recombination events from imprecise and noisy data. However, there are many more aspects of the likelihood function that could be examined, beyond coarse parameters like recombination rate or effective populations size. Theoretical explanations of this apparent robustness, and fully characterising the properties of the likelihood on the output of tsinfer and other methods is an important avenue for future work. Similarly, while there are many potential applications to concrete inferential problems, the details are nontrivial and must be consigned to future work.

Our results, while preliminary, suggest that “full ARGs”, in which each recombination and common ancestor event is precisely estimated, contain a significant degree of redundancy in terms of capturing information about the generative process. It is possible, therefore, that explicitly estimating *all* such events is not necessary to sample ARGs under the SMC (as ARGweaver and SINGER do). A combined approach, in which the uncertainty around the ordering of events that are weakly informed by the data is encoded structurally in the graph and other forms of uncertainty captured by sampling from the posterior, may yield further improvements in scalability for statistically rigorous inference. Another exciting prospect for ARG inference is the development of “hybrid” inference methods, where heuristic methods such as tsinfer or ARG-Needle are used to infer the details of the recent past in huge datasets such as UK Biobank, passing over to a rigorous model-based approach for the ancient past. This is similar in spirit to the widely-used approach of simulating complex dynamics of the recent past using detailed but slow models and using faster coalescent simulations in the distant past when the assumptions required are more justified (Bhaskar et al., 2014; Kelleher et al., 2018; Haller et al., 2018; Nelson et al., 2020). Just as forward and backward simulations are useful in different contexts and can be combined to great effect, large-scale heuristic and Bayesian sampling inference methods have their applications, and there may be synergy in their combination. The practical details of “handing over” between methods will require cooperation and precise communication; the tskit library, with its well-defined data model and flexible metadata support, is the ideal basis for this.

## Data availability

All scripts and data used for this manuscript are available at https://github.com/gertjanbisschop/smc-bit-paper, and an implementation of the algorithm is available at https://github.com/gertjanbisschop/runsmc.

## Acknowledgements

JK acknowledges EPSRC (research grant EP/X024881/1) and the Robertson Foundation. JK and PR acknowledge support from NIH (research grants HG011395 and HG012473).

## Appendix Algorithm

In this section we describe how to efficiently compute the likelihood under the SMC from an ARG stored using the `tskit` tree sequence encoding. The tree sequence format encodes relationships using *edges*, along with additional information that allows us to sequentially recover (and reason about their differences) efficiently (Kelleher et al., 2016; Ralph et al., 2020). The order of the edge deletions and insertions required to move from one local tree *T*_1_ to the next, *T*_2_ imposes the strict total ordering on edges (see definition (1)) that is needed to efficiently maintain the state of *I*_*e*_(*t*) for each focal edge *e*. Indeed, following the removal of edges that do not persist in *T*_2_, we simply declare that each subsequently added edge can only coalesce with the already-visited edges that make up the (partially) reconstructed local tree *T*_2_.

**Figure A1:**
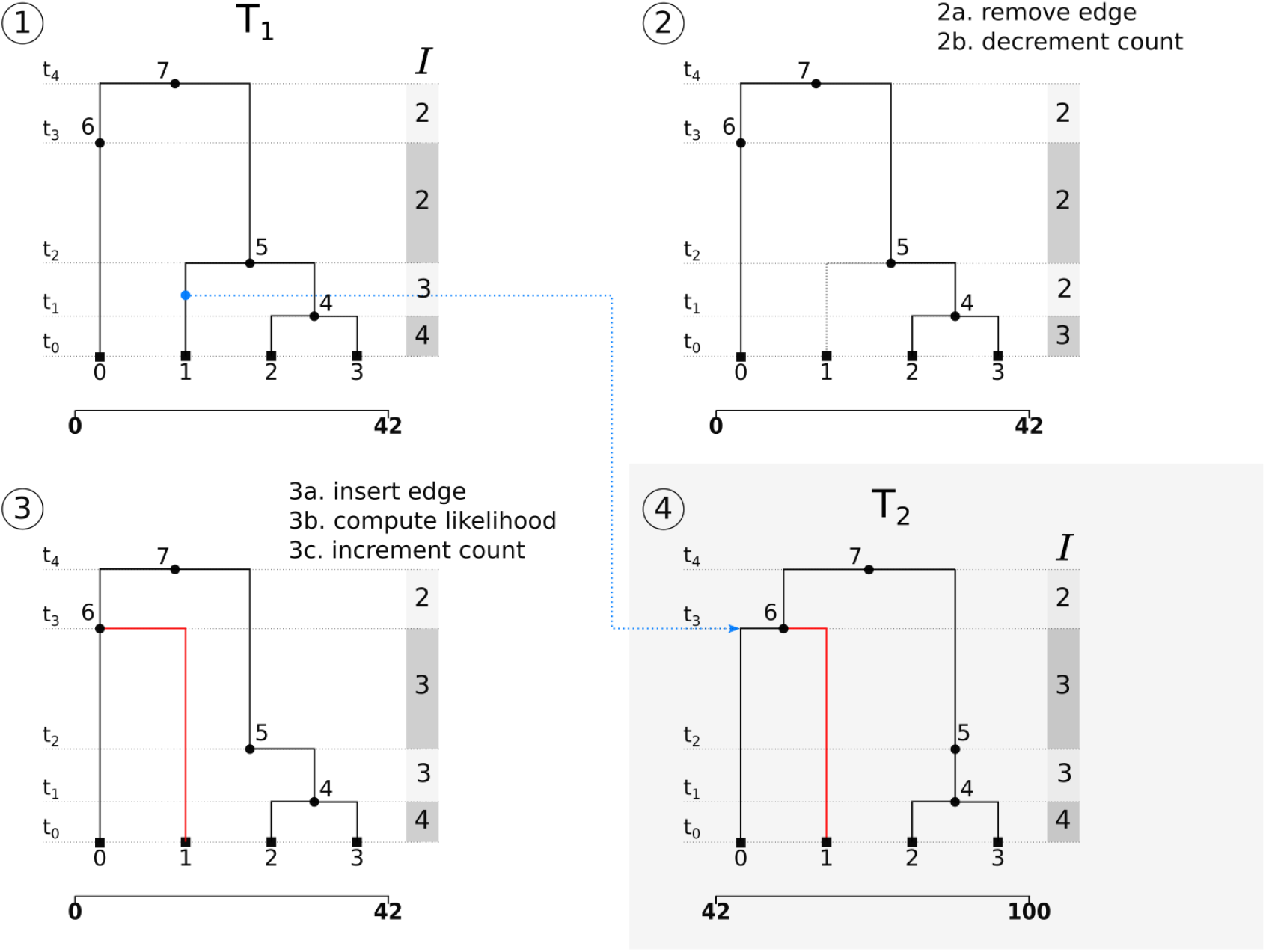
Likelihood computation algorithm. Moving along the genome we transition from local tree *T*_1_ to the next, *T*_2_ in three steps. Edges that do not persist in *T*_2_ are removed first. Edges starting at the left coordinate of *T*_2_ are inserted. After deletion or insertion of an edge the count vector *I* is updated along the corresponding internode intervals. Following an insertion, and prior to updating *I*, the likelihood for that edge is computed as *I* then represents the total number of lineages the lineage associated with this new edge can coalesce with. By moving along the genome in this way, each edge gets visited exactly once.

This allows us to compute a likelihood in a single pass over all edges. More concretely, the likelihood is computed by maintaining a single vector *I* that tracks the number of extant lineages in each internode interval (see Fig. A1). When moving from *T*_1_ to *T*_2_, the vector *I* is first decremented along the internode intervals spanned by the time of the child and the parent of all edges that do not persist in *T*_2_. For each subsequently inserted edge, we first compute the likelihood contribution of that edge by computing the integral in Eq. (5) using *I*, and then update the counts in *I*. We further keep track of the last parent a child node was connected to to detect recombination events.

